# Internal Validation of the ForenSeq Kintelligence Kit for Application to Forensic Genetic Genealogy

**DOI:** 10.1101/2022.10.28.514056

**Authors:** Michelle A. Peck, Alexander F. Koeppel, Erin M. Gorden, Jessica L. Bouchet, Mary C. Heaton, David A. Russell, Carmen R. Reedy, Christina M. Neal, Stephen D. Turner

## Abstract

Forensic Genetic Genealogy (FGG) requires high density single nucleotide polymorphism (SNP) profiles to infer distant relationships. The ForenSeq Kintelligence kit is a recently developed method targeting approximately 10,000 SNPs that were selected to be compatible with genetic genealogy databases, while avoiding medically relevant SNPs. The targeted PCR method is ideal for low input and degraded samples, which are particularly challenging for microarray analysis. We made modifications to library preparation and sample pooling to further enhance the performance of low input samples. To align this FGG method with forensic laboratory standards, we performed an internal validation study in accordance with the Scientific Working Group on DNA Analysis Methods Validation Guidelines for DNA Analysis Methods. The sensitivity and precision and accuracy studies produced accurate and reliable profiles down to 0.05 ng of DNA. The mixture and contamination studies demonstrated that the method could detect the presence of a minor contributor down to 0.02 ng, while system wide contamination was negligible. Nonprobative bone, fired shell casing, and adhesive tape samples performed well with inputs ranging from 0.05 to 1.0 ng of DNA. A bone with a high degradation index and tested at 0.1 ng DNA input resulted in a call rate of 92.9% and a heterozygosity rate of 0.387. We demonstrated the importance of both call rate and heterozygosity rate to assess profile performance. Validating FGG methods is critical to ensuring the reliability and utility of SNP profiles that will be uploaded to genetic genealogy databases for the purpose of generating investigative leads.

## Introduction

The power of high-density single nucleotide polymorphism (SNP) profiles to infer distant relationships to generate investigative leads in criminal casework and human identification efforts is rapidly advancing the forensic genomics field.^1,2^ Investigators have leveraged publicly available databases to solve cases that were previously unresolvable with STR technology.^2–6^ Microarray analysis and whole genome sequencing (WGS) are the most common approaches to generate the hundreds of thousands of SNPs necessary to perform Forensic Genetic Genealogy (FGG). Microarray analysis is a powerful tool but is limited by the high DNA input requirements and difficulty with degraded samples. While success with lower inputs has been demonstrated with microarray analysis, degradation remains a challenge, which can have a detrimental impact on kinship analysis.^7–10^ WGS and SNP capture have also been used in distant kinship analysis with particularly challenging samples.^11,12^ However, there are ethical concerns surrounding FGG that cannot be ignored, such as privacy concerns with interrogating the whole genome.^3,13–15^

To address some of these limitations, Verogen developed the ForenSeq^®^ Kintelligence kit that targets 10,230 SNPs spanning the genome, maintaining compatibility with direct-to-consumer kits that largely populate the publicly available databases, while excluding medically relevant SNPs.^16^ The targeted sequencing method is well-suited for low quantity samples, and the total input can reach high quantities even for samples with low concentration due to the ability to add up to 25 μL of extract. Further, the kit is designed for degraded samples with an average amplicon size of <150 bp.

The high-density profiles generated with microarray analysis and WGS enable distant kinship matching by the DNA segment sharing approach.^1^ In contrast, kinship matching with profiles generated by the Kintelligence kit uses the One-to-Many Kinship tool within GEDmatch PRO™ due to the reduced number of SNPs. The One-to-Many Kinship tool uses a windowed kinship algorithm^16^ that modifies the principal component analysis approaches of PC-Relate^17^ and PC-AiR^18^ to estimate genetic relatedness. While distant kinship matching (out to 4^th^ degree) has high sensitivity and specificity with this method, reduced call rates and loss of heterozygosity will diminish this performance.^16^

While FGG methodology is not currently held to the same forensic standards as other forensic DNA analyses, proper characterization and validation is critical to ensure the accuracy and utility of profiles generated for FGG investigations.^19^ Available performance data for the Kintelligence method includes the developmental validation of the MiSeq FGx™ Forensic Genomics System,^20^ which encompasses the ForenSeq™ library preparation method, MiSeq FGx™ sequencing, and the ForenSeq™ Universal Analysis Software methods that are the backbone of the Kintelligence kit, and a manufacturer datasheet and application notes.^21–23^ In accordance with the Scientific Working Group on DNA Analysis Methods (SWGDAM) Validation Guidelines for DNA Analysis Methods,^24^ we performed an internal validation of the Kintelligence kit, taking into consideration NGS-specific criteria. In this paper we present our internal validation of the Kintelligence kit performing sensitivity, precision and accuracy, mixture, contamination assessment, and nonprobative studies. We present protocol modifications that were used throughout the validation studies to improve the performance of low input samples.

## Materials and Methods

### Samples

The known control samples NA24385 (HG002), included as the kit positive (Verogen, San Diego, CA), and NA24143 (HG004), source NIST RM 8392 (National Institute of Standards and Technology, Gaithersburg, MD), were used throughout the testing with a range of inputs. The following DNA sample was obtained from the NIGMS Human Genetic Cell Repository at the Coriell Institute for Medical Research (Camden, NJ): NA24631. Quantity was confirmed of all controls with the Investigator^®^ Quantiplex^®^ Pro kit (QIAGEN, Hilden, Germany) on the ABI 7500 (Thermo Fisher Scientific, Waltham, MA) and dilutions were based on these results. Mixtures were created using control samples NA24631 (HG005) and NA24143 to evaluate mixtures with male (minor): female (major) ratios and from donors with different ancestries. Based on concentrations, mixtures of 1:2, 1:5, and 1:20 were created.

Nonprobative sample types were bones, tape, and fired shell casings (Table S1). Bone samples were obtained from the Southeast Texas Applied Forensic Science Facility and were extracted by Sam Houston State University using a modified InnoXtract™ method (InnoGenomics, New Orleans, LA) and PrepFiler™ BTA (Thermo Fisher Scientific). A wide range of bone types (i.e., femur, vertebra, tibia, and humerus), bone conditions (i.e., buried, burned, cremated, embalmed, and surface decomposition), and quantity and quality of extracts were tested. Two samples (both NA24385) from adhesive tape were extracted with the Tape Analysis For Forensic Identification (TAFFI) method as previously described.^25^ Two mock fired shell casings with DNA from anonymous donors were extracted with the Forensic Recovery of Identification from Shell Casings (FRISC) method as previously described.^26^ All extracts were quantified with the Investigator Quantiplex Pro kit on the ABI 7500, generating concentration values and degradation indices (DI).

### Library Preparation

The ForenSeq Kintelligence kit (Verogen) was used for library preparation with the following exceptions to the manufacturer’s reference guide.^27^ At the library purification step of the procedure, the final resuspension volume of RSB was reduced to 27 μL, with a transfer supernatant volume of 25 μL. Preliminary optimization experiments (data not shown) indicated that the decreased resuspension volume resulted in comparable and in some samples improved call rate over the standard resuspension volume. This was particularly important for boosting the library concentration of low input samples. Library quantification was performed with the Qubit™ dsDNA High Sensitivity Assay kit (Thermo Fisher Scientific) on the Qubit 4.0 (Thermo Fisher Scientific). Additionally, samples were run with the Agilent High Sensitivity DNA Kit (Agilent Technologies, Santa Clara, CA) on the Agilent 2100 Bioanalyzer system (Agilent Technologies) as a qualitative assessment.

Five separate library preparations were performed encompassing 58 libraries, with at least one negative control (NC) included with each preparation. The sensitivity study evaluated inputs of 1.0, 0.25, 0.1, and 0.05 ng in duplicate of NA24385 and NA24143 (N=16) (Figure S1). The precision and accuracy study evaluated inputs of 1.0 and 0.1 ng in duplicate of NA24385 and NA24143 performed by two different scientists (N=16). The replicates from the sensitivity study represented the results for Scientist 1. The mixture study evaluated the 1:2 and 1:5 mixture ratios at inputs of 1.0 and 0.1 ng in duplicate, and the 1:20 mixture ratio at 1.0 ng in duplicate (N=10). The nonprobative study evaluated seven extracts at 1.0 ng, four extracts ranging from approximately 0.25 to 0.5 ng, two extracts at 0.1 ng, and two extracts at approximately 0.05 ng (Table S1). The two samples at 0.1 ng were replicates of bone samples that were also processed at 1.0 ng. In total there were six NCs prepared.

### Sequencing

Library pooling was modified to use a molarity-based approach and the following guidelines for pool composition. Libraries with a sample input of 1.0 or 0.25 ng were pooled together in 4-plex sample pools, while libraries with a sample input of 0.1 or 0.05 ng were pooled together in 2-plex sample pools. NCs were only run in the 4-plex pools with high input libraries and did not count towards the sample count. To address NGS-specific sequencing sensitivity, a limited evaluation was performed sequencing the NA24385 sensitivity libraries at the opposite plexity as described above (i.e., 1.0 ng libraries in a 2-plex pool). The library volume used for pooling was based on the Qubit concentration and determined by the other libraries to achieve equal DNA input. Dilution of the denatured library pool was also adjusted to target 12 pM based on the estimated molarity. If denatured library pools were estimated to be less than 12 pM, no further dilution was performed prior to sequencing. Sequencing was performed on the MiSeq FGx (Verogen) according to the manufacturer’s MiSeq FGx Sequencing System reference guide.^28^ In total, 22 sequencing runs were performed consisting of 80 sequenced libraries (Figure S1). Ten sequencing runs were 2-plex pools and twelve were 4-plex pools. This included two 4-plex runs repeated by a second scientist for sequencing reproducibility. While the majority of the nonprobative samples were above 0.25 ng input and thus were sequenced in a 4-plex pool, due to sequencing constraints the low input samples were also sequenced in a 4-plex pool.

### Primary and Secondary Analysis

Primary analysis to call genotypes was automatically performed in the Universal Analysis Software (UAS).^29^ Two analysis methods were initially applied to samples in the sensitivity study to evaluate different read thresholds. The default manufacturer’s settings imposed an allelic >20X read count threshold, while the modified settings imposed an allelic >10X read count threshold. All other studies used the modified settings of 10X. No edits were made in the UAS. The sample reports containing genotype calls were exported from the UAS and secondary analysis to assess the studies was performed with R.^30^

To assess the accuracy of the genotype calls, the published genomes of NA24385 and NA24143 from the Genome in a Bottle Consortium (GIAB) were downloaded and filtered for the SNPs in the Kintelligence kit.^31^ These profiles only contain high confidence regions, which resulted in 9,375 and 9,353 SNPs for comparison with NA24385 and NA24143 Kintelligence profiles, respectively. Additionally, the published genomes from the 1000 Genomes Project were downloaded and filtered for the SNPs in the Kintelligence kit.^32^ The overall heterozygosity rate across these filtered genomes was computed, which served as a baseline for evaluating expected sample heterozygosity in several downstream analyses.

### GEDmatch PRO

To evaluate the performance of the Kintelligence kit with distant kinship matching, GEDmatch PRO (Verogen) reports were exported from the UAS for the sensitivity (10X and 20X thresholds) and nonprobative (10X) studies, excluding the fired shell casing samples. Samples were uploaded as target testing kits to GEDmatch PRO with permission from Verogen. The One-to-Many Kinship tool was applied to all samples and both the high confidence matches and expanded match list reports were downloaded. No attempt at re-identification of the samples was made.

## Results

### Library Preparation and Sequencing Performance

We generated 58 libraries across all studies with concentrations ranging from 0.13 ng/μL (0.1 ng input sample) to 4.9 ng/μL (1.0 ng input sample). Performance generally decreased with DNA input, yet there was substantial variability in library concentration within samples of the same DNA inputs (Table S2). Notably the NCs had a mean library concentration of 0.46 ng/μL, while the 0.05 ng/μL samples had a mean of only 0.21 ng/μL. However, Bioanalyzer traces were distinct between samples and NCs (Figure S2), suggesting that the concentrations observed in NCs were the result of primer dimer.

All 22 sequencing runs we performed were successful, passing UAS quality metrics and generating 6 – 35 million analyzed reads. The total number of reads analyzed per pool increased with increasing molarity of the pool (Figure S3), impacted in part by the initial DNA input of the libraries that comprised the sequencing pools. Similarly, the total number of reads analyzed per sample trended with the sample library molarity but was also impacted by the initial DNA input (Figure S4).

### Sensitivity

We evaluated the sensitivity of 16 high quality control samples by measuring the call rates at four DNA input levels (1.0, 0.25, 0.1, and 0.05 ng) using two different coverage thresholds for analysis (10X and 20X coverage). Call rate was greater than 95% for nearly all input levels at both analysis thresholds (Table 1). The only exceptions were the replicates of 0.05 ng at the 20X threshold (Figure 1A, Table S2). As expected, the mean call rate per input was always higher with the 10X threshold compared to the 20X threshold. The difference in mean call rate between analysis methods was statistically significant (t-test; p-values adjusted for multiple comparisons using the Benjamini-Hochberg false discovery rate (FDR))^33^ for the 0.1 ng (adjusted p<0.01) and 0.05 ng inputs (adjusted p<0.00001).

**Table 1.** Sensitivity study call rate, heterozygosity rate, and concordance rate mean results per DNA input and analysis threshold (n=4 per condition). Call rate was calculated with all 10,230 SNPs across individual samples. Heterozygosity rate was calculated with called SNPs across individual samples. Concordance rate was calculated with 9,375 SNPs and 9,353 SNPs for NA24385 and NA24143, respectively, based on high confidence regions in the GIAB profiles, across individual samples.

| **DNA Input**<br>**(ng)** | **Analysis**<br>**Threshold** | **Call Rate**<br>**Mean** | **Heterozygosity**<br>**Rate Mean** | **Concordance Rate**<br>**Mean** |
| --- | --- | --- | --- | --- |
| 0.05 | 10X | 0.962 | 0.409 | 0.944 |
| 0.05 | 20X | 0.934 | 0.383 | 0.921 |
| 0.1 | 10X | 0.981 | 0.441 | 0.976 |
| 0.1 | 20X | 0.966 | 0.428 | 0.965 |
| 0.25 | 10X | 0.989 | 0.453 | 0.989 |
| 0.25 | 20X | 0.978 | 0.444 | 0.982 |
| 1 | 10X | 0.987 | 0.455 | 0.992 |
| 1 | 20X | 0.975 | 0.449 | 0.988 |

**Figure 1.**
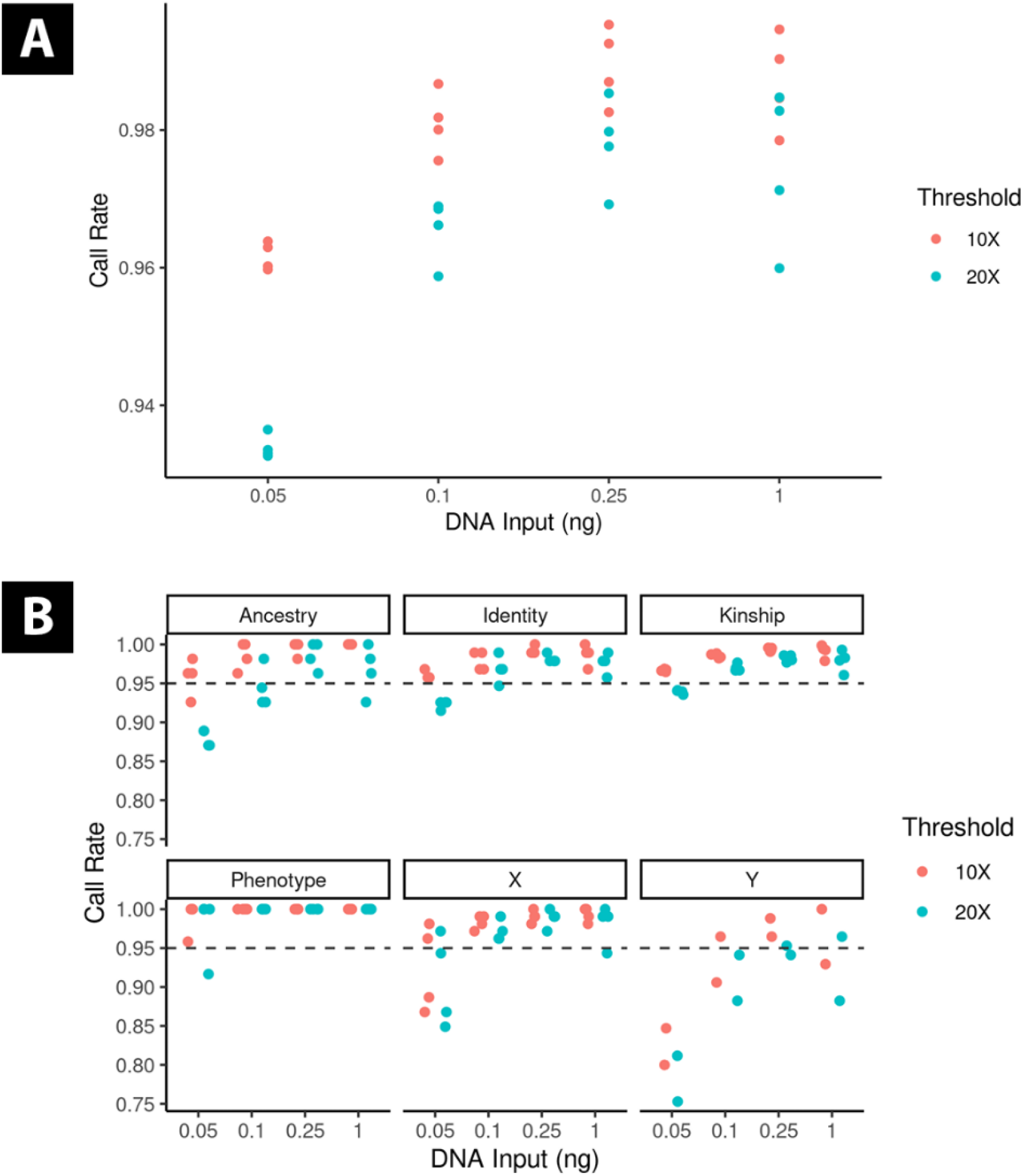
A) Sensitivity study call rate per sample for both 10X and 20X analysis methods. Call rates were higher with the 10X threshold and decreased with DNA input. B) Sensitivity study call rate by SNP category per sample for both 10X and 20X analysis methods. The dashed line represents 95% call rate. The Y-SNP call rates are only for the male sample NA24385. The inputs are discrete values, but the data points are slightly horizontally shifted (jittered) for visibility.

We next evaluated the call rates for specific subsets of SNPs, categorized based on their analytical purpose as annotated on the Kintelligence panel (e.g., ancestry, kinship). The call rates for each SNP category were generally consistent with the overall call rates per input (Figure 1B). Notably, the phenotype SNPs had a call rate of 100% for all but one sample, a replicate of NA24143 at the 0.05 ng input at both thresholds. The Y-SNPs were the worst performing type, dropping below an 85% call rate for all 0.05 ng replicates (data only shown for the male sample NA24385). While the call rate of the X-SNPs was high for most inputs and conditions, both replicates of the male sample NA24385 at 0.05 ng were notably lower (<90%). This may be a low input stochastic effect or could be related to the sex of the sample.

As a quality assurance check, we evaluated the heterozygosity rate for all samples, and compared them against a baseline rate derived from the 1000 Genomes Project samples. Across all populations in 1000 Genomes, heterozygosity averaged 0.431 ± 0.038 (three standard deviation range: 0.316 – 0.546). The observed heterozygosity for all inputs at both analysis thresholds fell within this range (Table 1).

Concordance compared to the known GIAB profiles was greater than 98% for 1.0 and 0.25 ng input samples (Table 1). Consistent with call rate, concordance decreased with lower inputs and was lower with the 20X threshold, however there was no significant difference between thresholds (t-test with FDR correction as described above). The lowest concordance rate was 92.1% with the 0.05 ng samples at 20X but was 94.4% at the 10X threshold. Most discordant calls (97.65%) were false homozygous calls due to allelic dropout, with higher levels observed in the 20X samples within each input level (Figure 2). Conversely, higher levels of false heterozygous calls (drop-in) were observed in the 10X samples. This is consistent with a higher threshold being more likely to miss a true second allele just below threshold, and a lower threshold being more likely to include a false allele just above threshold. We evaluated an intralocus balance (ILB) as a means to remove these false heterozygous calls, but determined it had minimal utility (Figure S5 and Table S3). Based on the sensitivity analysis, all other studies used the 10X threshold because it maximized SNP recovery and did not apply an ILB threshold.

**Figure 2.**
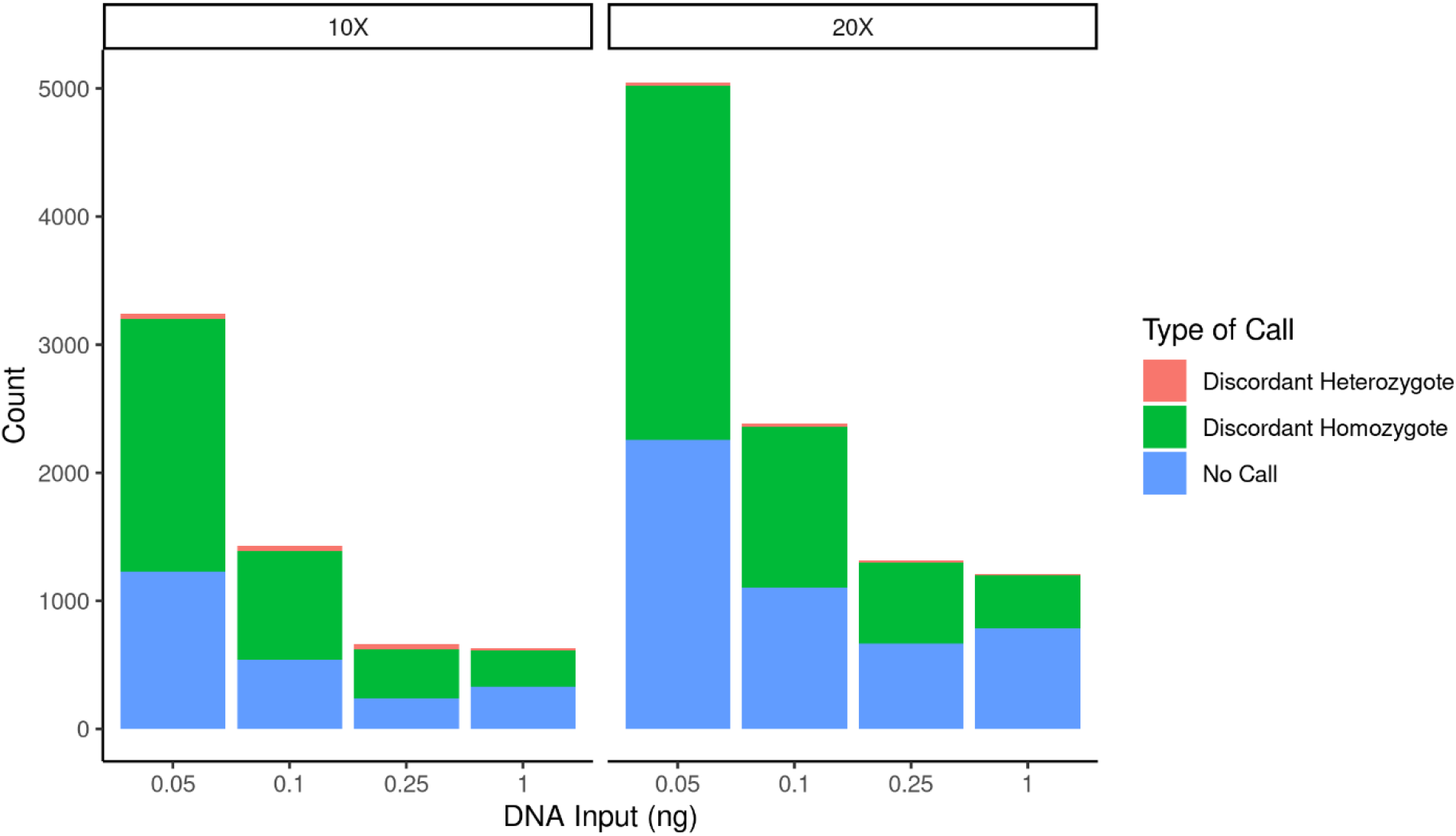
In the sensitivity study concordance evaluation, all inputs and analysis methods had high concordance to the known GIAB profiles. Displayed is the breakdown of counts for discordant heterozygotes (drop-in), discordant homozygotes (drop-out), and no call SNPs split by the analysis methods. Concordant SNPs counts are not shown due to extremely high values. In total, each input and analysis method evaluated 37,456 SNPs.

We additionally assessed sequencing sensitivity by evaluating the NA24385 libraries in both 2-plex and 4-plex pools for all inputs. There was minimal difference in call rate and concordance for the 1.0 and 0.25 ng samples (Table S4). However, both the call rate and concordance were approximately 1.8 and 3.7% higher for the 0.1 and 0.05 ng samples, respectively, when sequenced as a 2-plex pool.

### Precision and Accuracy

To assess the precision and accuracy of the assay, two independent scientists evaluated duplicate samples of NA24385 and NA24143 at 1.0 and 0.1 ng. The separate preparations resulted in different library concentrations but similar profile performance. The average library concentration for Scientist 2 was 4.2 and 2.1X higher at inputs of 0.1 and 1.0 ng, respectively (Figure S6). Yet, call rates were highly consistent for all samples with call rates >98.0% for all but one 0.1 ng sample and one 1.0 ng sample (Table S5). The accuracy of the profiles was established in the sensitivity study with Scientist 1 replicates compared against the GIAB known profiles. We assessed the precision of the assay by directly comparing profiles against the corresponding sample and input within scientist replicates (N=8 comparisons) for repeatability, and between scientist replicates (N=16) for reproducibility. The mean concordance rates for both repeatability and reproducibly were greater than 96.9 and 99.1% for 0.1 and 1.0 ng, respectively (Figure 3).

**Figure 3.**
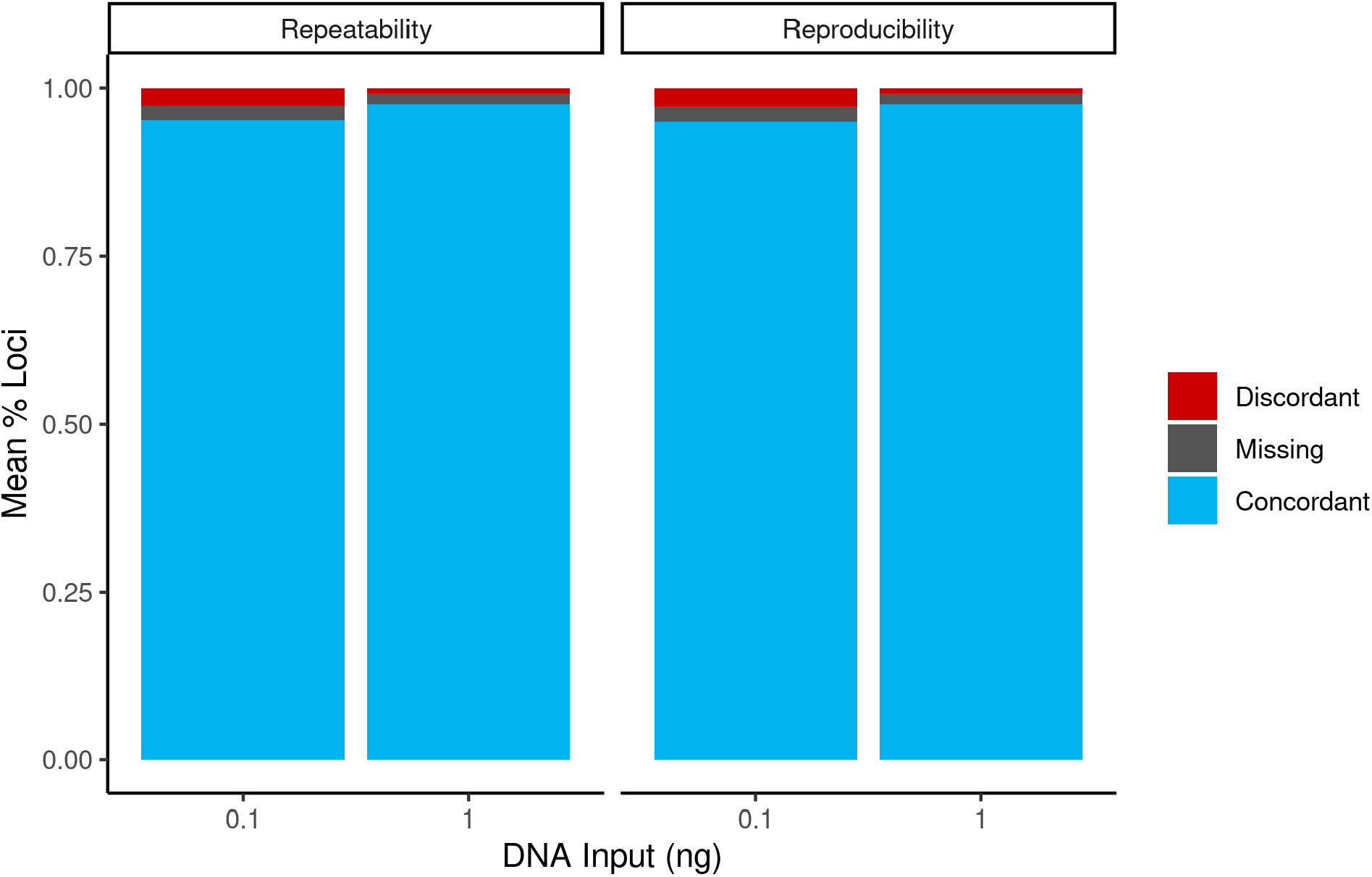
Percentage of SNP loci that were concordant, discordant, and missing when comparing replicate profiles within a scientist for repeatability and between scientists for reproducibility. The concordance rate for both assessments were >96.9 and >99.1% for 0.1 and 1.0 ng, respectively.

Additionally, we repeated the sequencing of two 4-plex pools consisting of 1.0 ng libraries and assessed the concordance between the specific replicate libraries (N=8). The concordance averaged 99.4% demonstrating sequencing reproducibility.

### Mixtures

We evaluated mixture ratios of 1:2, 1:5, and 1:20 at DNA inputs of 1.0 and 0.1 ng (excluding the 1:20 mixture). Library and sequencing performance of the mixtures was consistent with single-source samples but were flagged as mixtures within the UAS. The call rate mean was 97.4% for the 0.1 ng samples and 99.6% for the 1.0 ng samples (Table S6). The distinguishing feature of these samples was the elevated heterozygosity rate. All inputs and mixture ratios had >0.60 heterozygosity rate, notably 0.795 for the 1:2 samples at 1.0 ng, well above the upper range of 0.546 established from the 1000 Genomes dataset (Figure 4A). Comparing the mixture profiles to a single-source NA24143 sample (the major component of all mixtures), concordance averaged <81.3%, indicating the detection of the minor contributor in all samples (Figure 4B).

**Figure 4.**
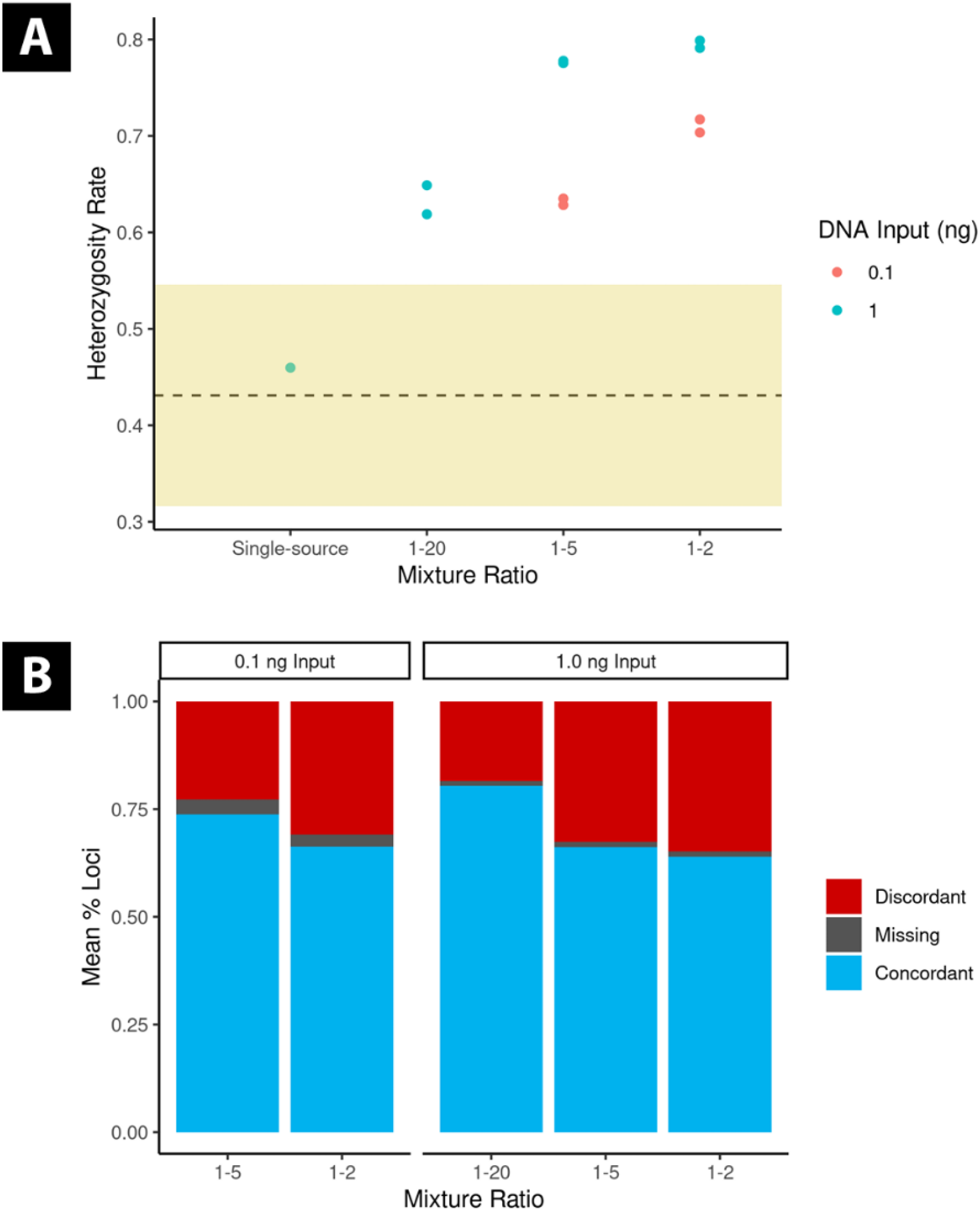
A) Heterozygosity rate for the mixture samples compared to a single-source NA24143 sample, with the range from the 1000 Genomes samples highlighted. The dashed line represents the mean of 1000 Genomes heterozygosity rate (0.431), and the yellow band indicates three standard deviations above and below the mean (0.316 – 0.546). All mixture ratios at both inputs were well above this range. B) The percentage of SNP loci that were concordant, discordant, and missing when comparing the mixture samples to a single source NA24143 sample (major component of all mixtures) are displayed. High rates of discordance indicate that the minor profile was detected in all samples.

### Contamination Assessment

During the course of the validation studies, we prepared six NC libraries and performed twelve sequencing events of these libraries. The number of SNPs called was drastically lower than what was observed in all other samples, ranging from 7 to 92 called SNPs (<1.0% call rate) (Table S7). All called SNPs were homozygous genotypes. We chose to not sequence NCs with low input samples, specifically in 2-plex sequencing pools, to avoid further diluting the samples. As a simple check to compare performance, the sequencing sensitivity evaluation included two NCs in a 2-plex sequencing pool. The number of SNPs called only increased from 13 to 32 and 7 to 15 and were all homozygous genotypes.

### Nonprobative Samples

The nonprobative study focused on challenging bone samples from a range of bone types and insults. Of the eleven bones tested at inputs ranging from 0.1 to 1.0 ng, seven (63.6%) had call rates ≥96.9% (Figure 5 and Table S8). Bone 41 was a cremated bone and had the lowest call rate at 92.3% of the 1.0 ng input samples but was still sufficiently high. However, when this extract was tested at 0.1 ng, the call rate was drastically reduced to 63.4% with a heterozygosity rate of only 0.174, well below the lower end of the expected range (0.316). Two extracts of the same bone (Bone 44 and Bone 46) tested at 1.0 ng DNA input, each had high degradation indices at 14.3 and 26.1, respectively, but resulted in high call rates of 97.7% and 96.9%. Further, when the same extract as Bone 44 was tested using 0.1 ng of DNA input (Bone 45) the call rate decreased minimally to 92.9% with a heterozygosity rate within range. Bone 48 also had a high degradation index at 32.4 and had one of the lowest call rates at 79.6%, indicating some sample-dependent performance. Yet, the heterozygosity rate of this sample was within range at 0.387.

**Figure 5.**
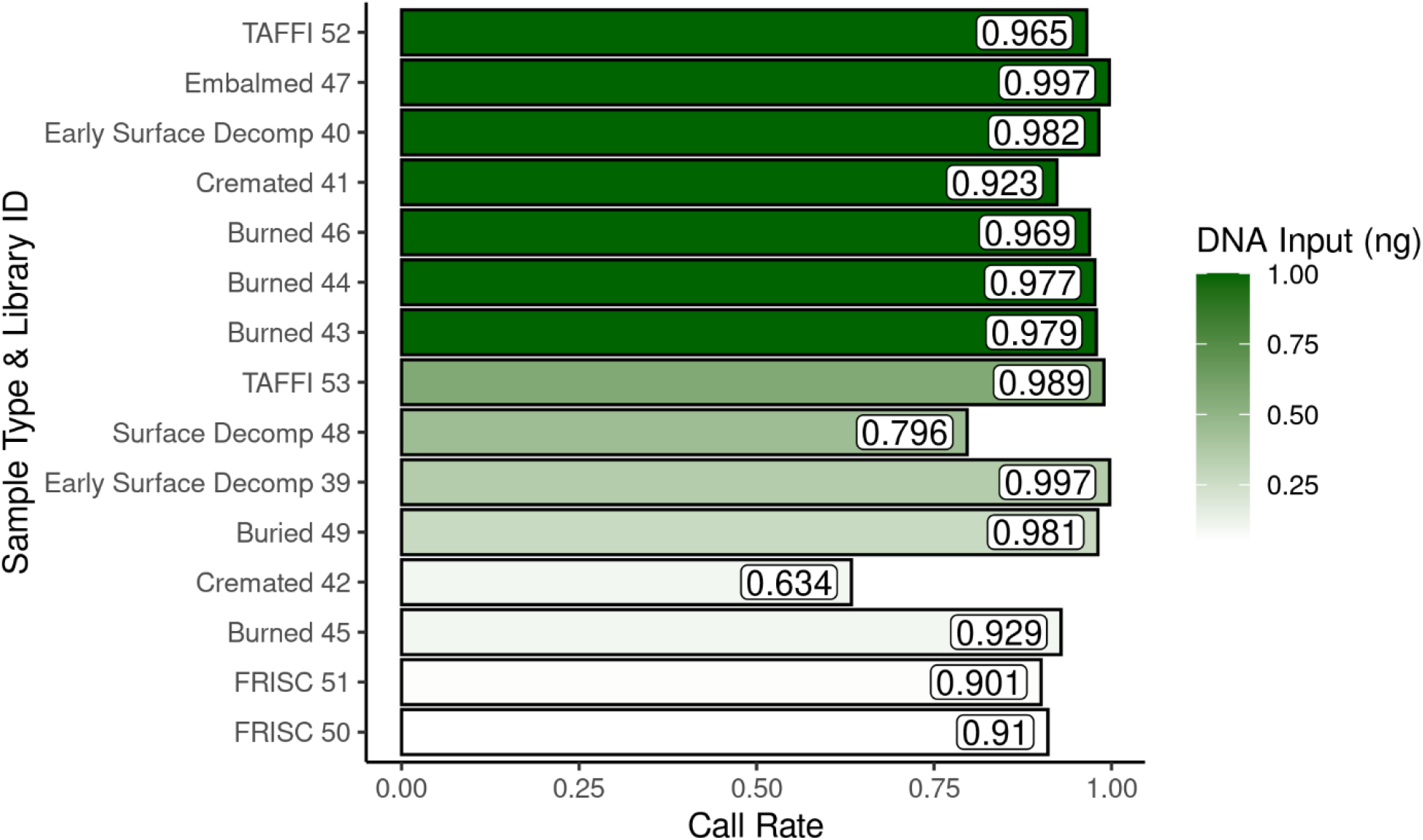
Nonprobative call rate ordered by descending DNA input. The following samples were the same extract tested at different inputs: Cremated 41 & Cremated 42, and Burned 44 & Burned 45.

Additionally, we evaluated two adhesive tape and two fired shell casing samples. The adhesive tape samples were tested at 1.0 and 0.569 ng resulting in high call rates of 96.5 and 98.9%, respectively. The fired shell casing extracts were tested at low inputs of 0.052 and 0.062 ng, which both resulted in call rates of >90%. Notably, the nonprobative samples were all sequenced at 4-plex sample pools, due to sequencing constraints. In practice, the low input samples (<0.25 ng) would be sequenced in 2-plex pools, likely resulting in higher call rates.

### GEDmatch PRO

As a proof of principle, we evaluated the NA24385 samples at 10X and 20X from the sensitivity study along with the nonprobative samples at 10X with the One-to-Many Kinship tool in GEDmatch PRO. Consistent match results to a likely 4^th^ degree relative were observed with the NA24385 samples across inputs and analysis methods, with the exception of one 0.05 ng replicate at 20X not detecting the match (Figure 6). Notably, the replicate that did not match had the lowest heterozygosity rate of 0.362. There was some variability in the relationship degree assignment (5^th^ degree for 13/15 replicates) and high confidence matching (13/15 replicates), as categorized by the tool. Similarly, the match results of nonprobative samples with replicates available for comparison were also consistent across inputs (Table 2). The TAFFI samples (donor NA24385) correctly matched to the same 4^th^ degree relative as the sensitivity samples, albeit also at a 5^th^ degree relationship, demonstrating the same detection with high-quality and nonprobative samples. The same high confidence second degree match resulted with Bone D for two replicates at 1.0 ng and one replicate at 0.1 ng. Bone A also had a consistent match between two replicates in the expanded match list, though one replicate had several other matches with similar results.

**Figure 6.**
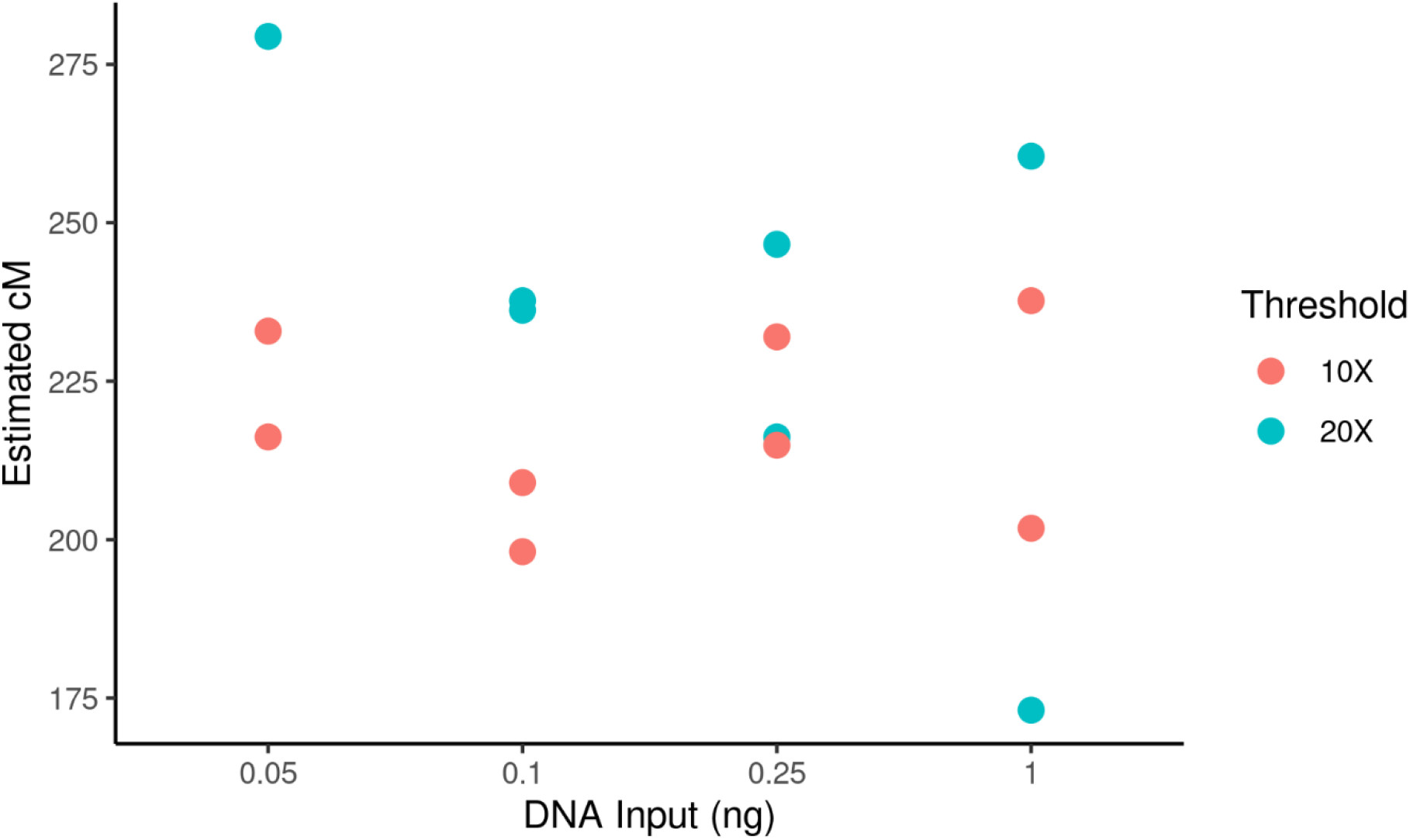
One-To-Many Kinship tool match results to a likely 4th degree relative for the NA24385 samples from the sensitivity study. One of the 0.05 ng replicates at 20X did not match to this individual and therefore is not displayed. The two highest cM values (0.05 ng at 20X and 1.0 ng at 20X) were flagged as 4^th^ degree while the rest were 5^th^ degree. All matches were high confidence except for the two lowest cM values (1.0 ng at 20X and 0.1 ng at 10X).

**Table 2.** GEDmatch PRO One-to-Many results for nonprobative samples with replicates available to compare. The names of the match results were anonymized. The match for the NA24385 TAFFI samples is the same relative as the matches for the sensitivity study (Figure 6). ^a^ The lower number of overlapping SNPs could be caused by the Alias Z profile being generated from an older microarray kit (Personal communication; Verogen, August 10, 2022).

| Donor | Library ID | Input (ng) | Call Rate | Match | Shared cM | Degree | High Confidence | Overlapping SNPs |
| --- | --- | --- | --- | --- | --- | --- | --- | --- |
| Bone D | Burned 44 | 1 | 0.977 | Alias X | 1333.9 | 2nd | yes | 9346 |
| Bone D | Burned 46 | 1 | 0.969 | Alias X | 1377.7 | 2nd | yes | 9279 |
| Bone D | Burned 45 | 0.1 | 0.929 | Alias X | 1136.6 | 2nd | yes | 8895 |
| Bone A | Early Surface Decomp 40 | 1 | 0.982 | Alias Y | 128.7 | 5th | no | 9629 |
| Bone A | Early Surface Decomp 39 | 0.36 | 0.997 | Alias Y | 126.6 | 5th | no | 9780 |
| NA24385 | TAFFI 52 | 1 | 0.965 | Alias Z | 227.8 | 5th | yes | 7621<sup>a</sup> |
| NA24385 | TAFFI 53 | 0.57 | 0.989 | Alias Z | 220.3 | 5th | yes | 7803<sup>a</sup> |

## Discussion

The Kintelligence kit produced accurate and reliable data throughout the validation studies, satisfying the SWGDAM requirements for internal validation. The high-quality control samples in the sensitivity study demonstrated high call rates and concordant results compared to the known GIAB profiles down to 0.05 ng. In comparing the 10X and 20X thresholds (manufacturer’s recommendation), we determined higher call rates and higher concordance resulted with the 10X threshold. We also evaluated an ILB threshold, but determined it only minimally removed discordant SNPs while removing more concordant SNPs. Applying the 10X threshold maximizes the data retained and was sufficient for high input samples at a 4-plex pool and low input samples at a 2-plex pool. The sequencing sensitivity study confirmed comparable performance in call rate and concordance of the high input samples in a 2-plex and 4-plex pool. Processing high input samples in a 4-plex pool thus increases throughput and reduces sequencing cost. Further, consistent GEDmatch PRO match results were observed at all input levels.

Evaluating the performance of 1.0 and 0.1 ng samples prepared by two scientists, indicated that while library concentration can differ, the sequencing results were highly repeatable and reproducible. The molarity-based approach likely contributed to this balanced performance. We demonstrated the validity of the molarity-based pooling approach and its significant impact on low quantity samples. The assay was shown to be highly repeatable and reproducible at both the library preparation and sequencing level.

The mixture study and contamination studies established heterozygosity as the evaluation criteria which would preclude a sample from moving forward to kinship analysis. The minor contributor was detected in all mixtures evaluated as demonstrated by the low concordance rates with the major donor, which corresponded to elevated heterozygosity rates outside the expected range. The lowest minor contributor portion was approximately 0.02 ng in the 1:5 mixture at 0.1 ng input. Widespread contamination was never observed in the NCs and a 4-plex sequencing pool was sufficient for assessment. The NCs were characterized by low call rates (≤1.0%) and the detection of exclusively homozygous SNPs.

The Kintelligence kit was able to produce profiles with high call rates with DNA extracted from challenging bone samples, adhesive tape, and fired shell casings. These high call rates were observed with degraded samples and low inputs. Many of the bone samples were able to reach the 1.0 ng target input, as a result of the high-volume input into library preparation. Heterozygosity rate was also useful for assessing the quality of profiles in addition to the call rate. Our evaluation of Bone D in GEDmatch PRO resulted in consistent match results to a 2^nd^ degree relative, even with a replicate with diminished call rate that had a heterozygosity rate within range. Snedecor et al. demonstrated that the windowed kinship algorithm used by GEDmatch PRO is relatively robust to diminished numbers of SNPs called and loss of heterozygosity, but the impact depends on the degree of relationship.^16^ As is often the case with challenging case samples, stochastic drop-out is expected causing decreased call rate and loss of heterozygosity to be compounded. Thus, call rate and heterozygosity should be considered in conjunction with the degree of relation that is detected to help provide guidance on the quality of the match.

## Conclusions

We validated the ForenSeq Kintelligence kit for producing quality SNP profiles for application to FGG. Modifications to the library resuspension volume and library pooling approach were critical to enhancing the performance of low quantity samples. The Kintelligence kit demonstrated sensitivity down to 0.05 ng with high quality samples, producing accurate and reproducible results. Performance with nonprobative samples was robust across a range of sample types, sample quality, and inputs, most notably with bone samples. Call rate and heterozygosity should be used to inform the quality of the profile generated. The Kintelligence kit is another approach that can be used to generate SNP data for distant kinship inference from challenging forensic samples or where microarray analysis or WGS are not an option.

## Acknowledgments

The authors would like to thank Rachel Houston and Jennifer Snedeker from Sam Houston State University for providing the bone extracts for the nonprobative testing. We would like to acknowledge the Sam Houston State University Department of Forensic Science as well as the Southeast Texas Applied Forensic Science Facility and the donors and their loved ones, without whom this research would not be possible. We also would like to thank Curt Hewitt, Signature Science, LLC, for providing the fired shell casing and adhesive tape extracts. Finally, we would like to thank Meredith Turnbough and June Snedecor from Verogen for helpful discussions on Kintelligence and GEDmatch PRO.

## Author Contributions

Michelle A. Peck: Conceptualization, Methodology, Validation, Formal Analysis, Investigation, Data Curation, Writing – Original Draft, Writing – Review and Editing, Visualization

Alexander F. Koeppel: Validation, Formal Analysis, Data Curation, Writing – Original Draft, Writing – Review and Editing, Visualization

Erin M. Gorden: Methodology, Writing – Review and Editing

Jessica L. Bouchet: Validation, Investigation

Mary C. Heaton: Methodology, Writing – Review and Editing

David A. Russell: Methodology, Writing – Review and Editing

Carmen R. Reedy: Conceptualization, Methodology, Writing – Review and Editing, Supervision, Project Administration

Christina M. Neal: Conceptualization, Methodology, Investigation, Writing – Review and Editing, Supervision, Project Administration

Stephen D. Turner: Methodology, Validation, Formal Analysis, Data Curation, Writing – Original Draft, Writing – Review and Editing, Supervision, Project Administration

## Author Disclosure Statement

The authors M.A.P., A.F.K, E.M.G., M.C.H., D.A.R., C.R.R., C.M.N., and S.D.T. are employees of Signature Science, LLC and have no competing interests to disclose.

## Funding Statement

This work was fully funded by Signature Science, LLC.

## Notes

### Competing Interest Statement

The authors have declared no competing interest.

