## Supplemental Tables and Figures for "Internal Validation of the ForenSeq Kintelligence Kit for Application to Forensic Genetic Genealogy"

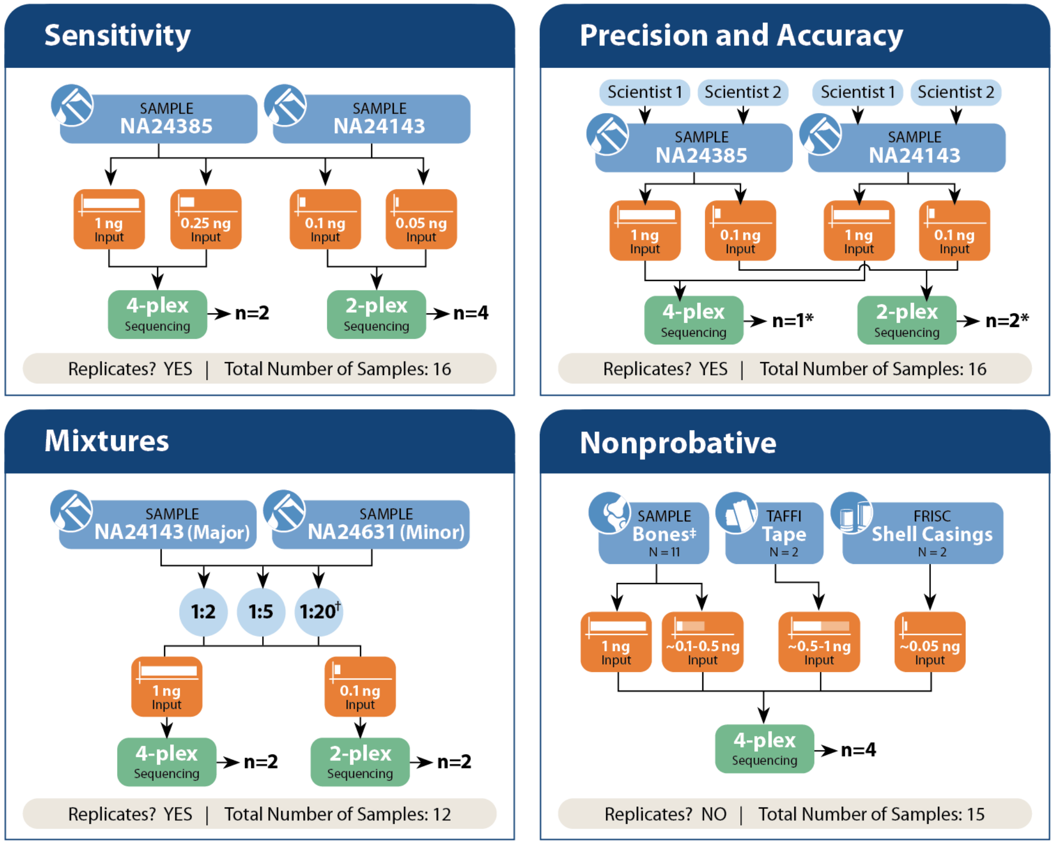


**Figure S1.** Overview of each study design. Studies not directly shown are sequencing reproducibility and sequencing sensitivity. Sequencing reproducibility repeated the sequencing of a 4-plex pool from sensitivity and from precision and accuracy, accounting for two additional 4-plex pools. Sequencing sensitivity tested the NA24385 libraries from sensitivity in two 2-plex pools and one 4-plex pool.
*The assessment of precision and accuracy used data from the sensitivity study from two 2-plex pools and one 4-plex pool.
†The 1:20 mixture ratio was not tested at 0.1 ng input.
‡Includes early surface decomposition, surface decomposition, buried, embalmed, burned, and cremated bones.

**Table S1.** Nonprobative sample details including extract and library information.

| | | | Extract Details | | | | | Library Details | | |
|---|---|---|---|---|---|---|---|---|---|---|
| Sample Type | Donor ID | Sample Details | Extract ID | Extraction Method | Concentration (ng/µL) | Degradation Index | IPC Value | Library ID | Input Volume (µL) | Total Input (ng) |
| Bone | A | femur, early surface decomp | 1 | InnoXtract | 0.0942 | 1.16 | 27.22 | 39 | 3.79 | 0.357 |
| Bone | A | femur, early surface decomp | 2 | PrepFiler BTA | 0.1150 | 1.22 | 26.93 | 40 | 8.70 | 1.000 |
| Bone | B | vertebra, cremated | 3 | InnoXtract | 0.0481 | 2.67 | 27.49 | 41 | 20.79 | 1.000 |
| Bone | B | vertebra, cremated | 3 | InnoXtract | 0.0481 | 2.67 | 27.49 | 42 | 2.08 | 0.100 |
| Bone | C | femur, burned | 4 | InnoXtract | 0.3638 | 9.92 | 26.90 | 43 | 2.75 | 1.000 |
| Bone | D | tibia, burned | 5 | InnoXtract | 0.3235 | 14.25 | 27.00 | 44 | 3.09 | 1.000 |
| Bone | D | tibia, burned | 5 | InnoXtract | 0.0500 | 14.25 | 27.00 | 45 | 2.00 | 0.100 |
| Bone | D | tibia, burned | 6 | PrepFiler BTA | 0.0420 | 26.08 | 27.98 | 46 | 23.81 | 1.000 |
| Bone | E | femur, embalmed | 7 | InnoXtract | 0.0981 | 3.69 | 27.02 | 47 | 10.19 | 1.000 |
| Bone | F | tibia, surface decomp | 8 | InnoXtract | 0.0185 | 32.40 | 26.94 | 48 | 24.00 | 0.444 |
| Bone | G | humerus & tibia, buried | 9 | InnoXtract | 0.0108 | 4.24 | 27.00 | 49 | 25.00 | 0.271 |
| Shell casing | H | -- | 10 | FRISC | 0.0045 | -- | -- | 50 | 11.5 | 0.052 |
| Shell casing | H | -- | 11 | FRISC | 0.0055 | -- | -- | 51 | 11.4 | 0.062 |
| Adhesive tape | NA24385 | -- | 12 | TAFFI | 0.0513 | -- | -- | 52 | 19.48 | 1.000 |
| Adhesive tape | NA24385 | -- | 13 | TAFFI | 0.0569 | -- | -- | 53 | 10.00 | 0.569 |

**Table S2.** Average library concentration values for all studies excluding the nonprobative study.

| DNA Input (ng) | Average Library Concentration (ng/µL) | Standard Deviation of Library Concentration | Library Count |
|---|---|---|---|
| 0 | 0.46 | 0.38 | 5 |
| 0.05 | 0.21 | 0.04 | 4 |
| 0.1 | 0.60 | 0.39 | 14 |
| 0.25 | 0.60 | 0.21 | 4 |
| 1 | 2.62 | 1.17 | 14 |

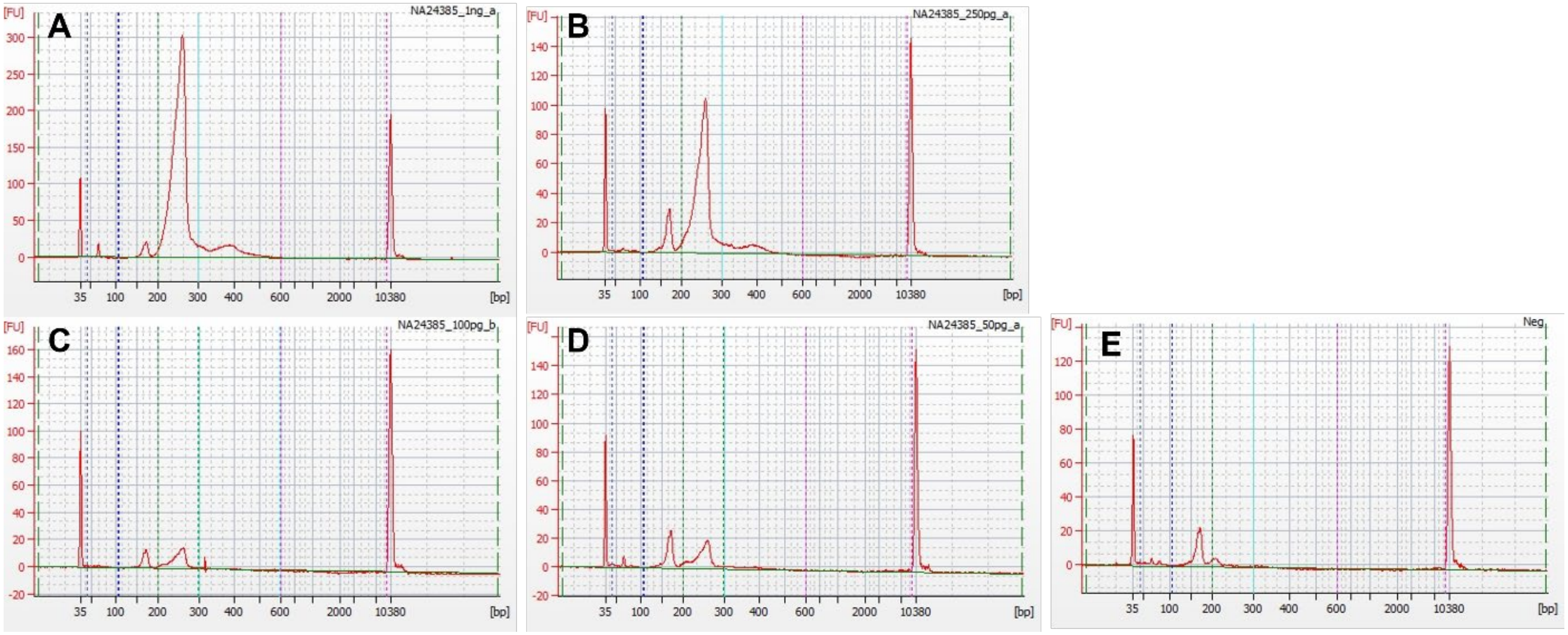


**Figure S2.** Representative bioanalyzer traces of NA24385 libraries at 1.0 ng (A), 0.25 ng (B), 0.1 ng (C), and 0.05 ng (D) and a negative control (E). The peak visible in the negative control and likely contributing to the higher library concentrations is consistent with primer dimer.

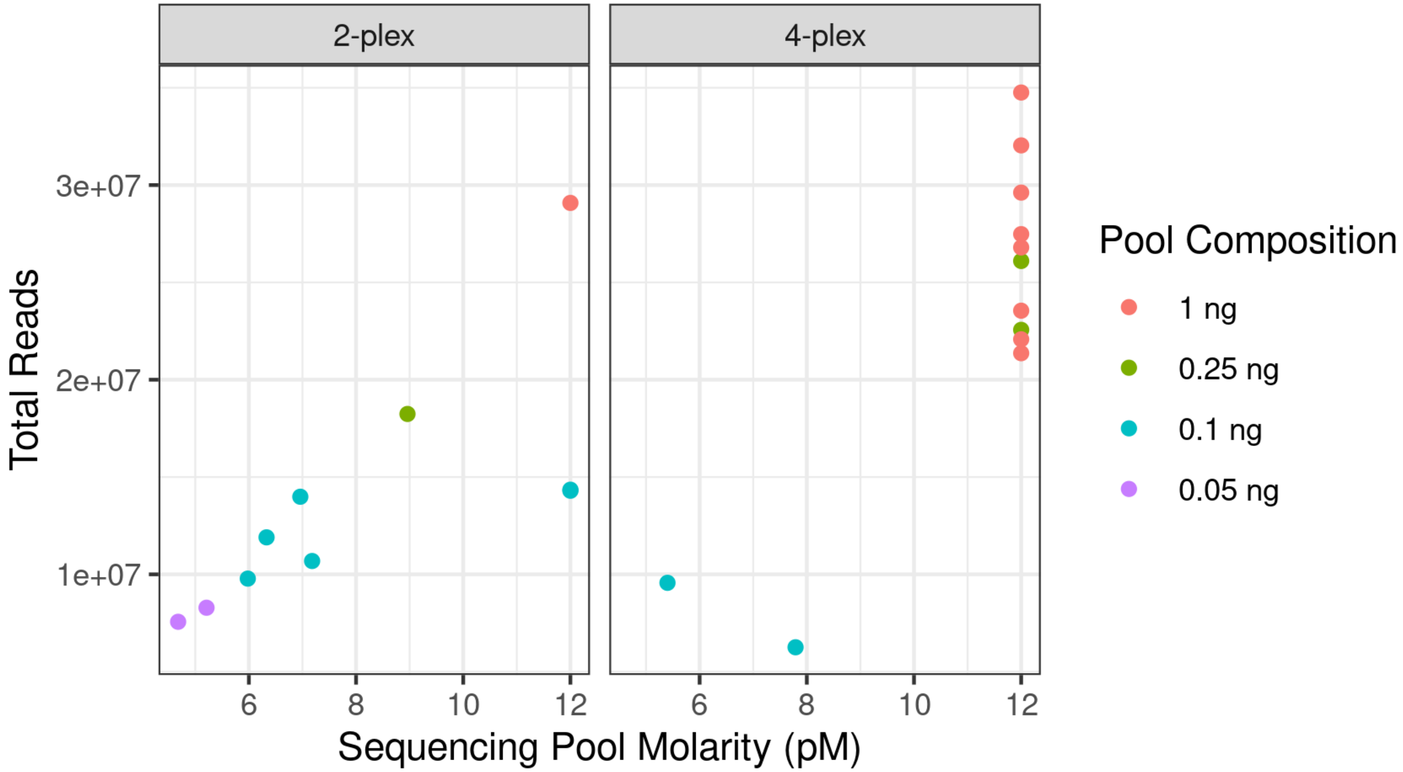


**Figure S3.** Total reads analyzed per sequencing run plotted against the estimated molarity of the sequencing pool for all sequencing runs split by the sample plexity. The colors indicate the inputs of the libraries that comprised the sequencing pools.

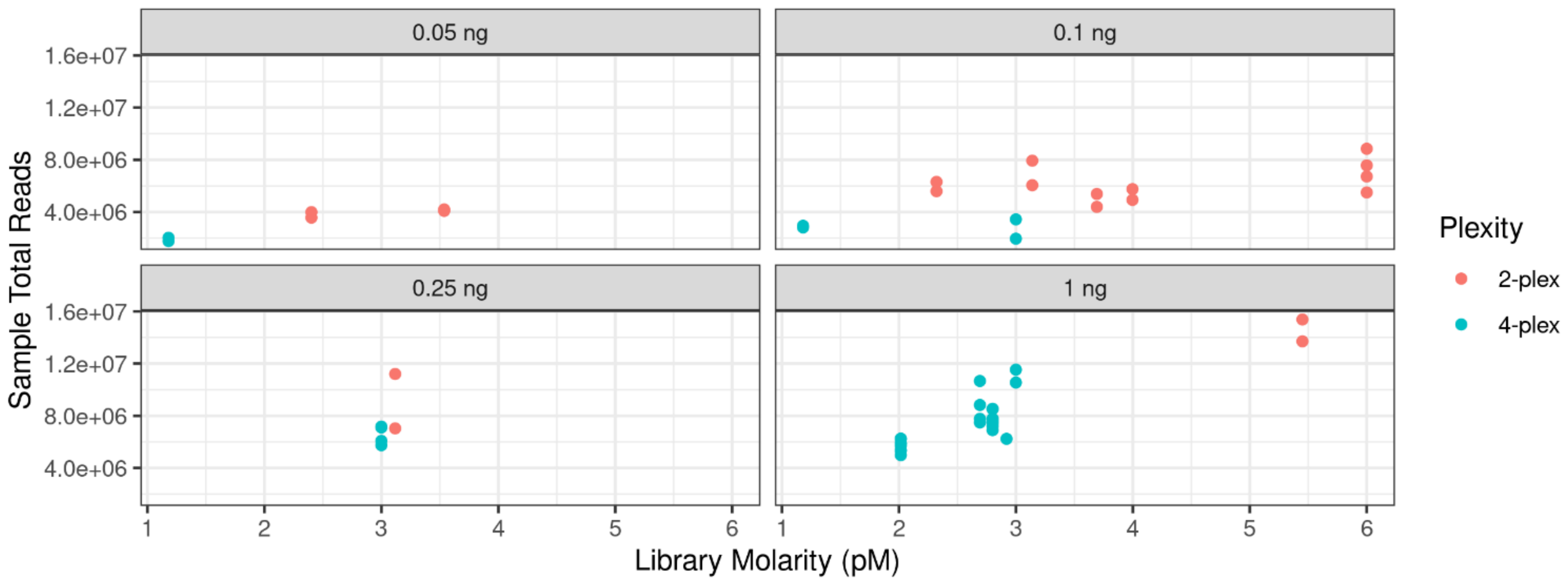


**Figure S4.** Total reads analyzed per sample plotted against the estimated molarity of the libraries for NA24385 and NA24143 split by the DNA input. The colors indicate the sequencing pool plexity.

**Table S2.** Sensitivity study sample profile results analyzed at both 10X and 20X thresholds, arranged by ascending DNA input (ng).

| Sample Name | Library ID | DNA Input (ng) | Analysis Threshold | Missed SNPs | Heterozygous SNPs | Called SNPs | Call Rate | Heterozygosity Rate | Intralocus Balance Mean |
|---|---|---|---|---|---|---|---|---|---|
| NA24143 | 33 | 0.05 | 10X | 407 | 4102 | 9823 | 0.960 | 0.418 | 0.561 |
| NA24143 | 33 | 0.05 | 20X | 685 | 3771 | 9545 | 0.933 | 0.395 | 0.578 |
| NA24143 | 37 | 0.05 | 10X | 412 | 4097 | 9818 | 0.960 | 0.417 | 0.555 |
| NA24143 | 37 | 0.05 | 20X | 689 | 3775 | 9541 | 0.933 | 0.396 | 0.571 |
| NA24385 | 24 | 0.05 | 10X | 370 | 4005 | 9860 | 0.964 | 0.406 | 0.532 |
| NA24385 | 24 | 0.05 | 20X | 650 | 3624 | 9580 | 0.936 | 0.378 | 0.553 |
| NA24385 | 28 | 0.05 | 10X | 379 | 3896 | 9851 | 0.963 | 0.395 | 0.515 |
| NA24385 | 28 | 0.05 | 20X | 680 | 3461 | 9550 | 0.934 | 0.362 | 0.537 |
| NA24143 | 30 | 0.1 | 10X | 250 | 4396 | 9980 | 0.976 | 0.440 | 0.627 |
| NA24143 | 30 | 0.1 | 20X | 422 | 4212 | 9808 | 0.959 | 0.429 | 0.638 |
| NA24143 | 32 | 0.1 | 10X | 204 | 4502 | 10026 | 0.980 | 0.449 | 0.635 |
| NA24143 | 32 | 0.1 | 20X | 322 | 4337 | 9908 | 0.969 | 0.438 | 0.644 |
| NA24385 | 21 | 0.1 | 10X | 136 | 4431 | 10094 | 0.987 | 0.439 | 0.602 |
| NA24385 | 21 | 0.1 | 20X | 318 | 4223 | 9912 | 0.969 | 0.426 | 0.613 |
| NA24385 | 23 | 0.1 | 10X | 186 | 4362 | 10044 | 0.982 | 0.434 | 0.597 |
| NA24385 | 23 | 0.1 | 20X | 346 | 4123 | 9884 | 0.966 | 0.417 | 0.610 |
| NA24143 | 34 | 0.25 | 10X | 178 | 4566 | 10052 | 0.983 | 0.454 | 0.74 |
| NA24143 | 34 | 0.25 | 20X | 315 | 4426 | 9915 | 0.969 | 0.446 | 0.748 |
| NA24143 | 36 | 0.25 | 10X | 133 | 4625 | 10097 | 0.987 | 0.458 | 0.733 |
| NA24143 | 36 | 0.25 | 20X | 229 | 4497 | 10001 | 0.978 | 0.450 | 0.739 |
| NA24385 | 25 | 0.25 | 10X | 48 | 4599 | 10182 | 0.995 | 0.452 | 0.706 |
| NA24385 | 25 | 0.25 | 20X | 150 | 4457 | 10080 | 0.985 | 0.442 | 0.712 |
| NA24385 | 27 | 0.25 | 10X | 76 | 4559 | 10154 | 0.993 | 0.449 | 0.705 |
| NA24385 | 27 | 0.25 | 20X | 207 | 4389 | 10023 | 0.980 | 0.438 | 0.714 |
| NA24143 | 31 | 1 | 10X | 157 | 4613 | 10073 | 0.985 | 0.458 | 0.813 |
| NA24143 | 31 | 1 | 20X | 294 | 4487 | 9936 | 0.971 | 0.452 | 0.817 |
| NA24143 | 38 | 1 | 10X | 99 | 4672 | 10131 | 0.990 | 0.461 | 0.817 |

| Sample Name | Library ID | DNA Input (ng) | Analysis Threshold | Missed SNPs | Heterozygous SNPs | Called SNPs | Call Rate | Heterozygosity Rate | Intralocus Balance Mean |
|---|---|---|---|---|---|---|---|---|---|
| NA24143 | 38 | 1 | 20X | 156 | 4593 | 10074 | 0.985 | 0.456 | 0.822 |
| NA24385 | 22 | 1 | 10X | 220 | 4498 | 10010 | 0.978 | 0.449 | 0.786 |
| NA24385 | 22 | 1 | 20X | 410 | 4335 | 9820 | 0.960 | 0.441 | 0.794 |
| NA24385 | 29 | 1 | 10X | 55 | 4599 | 10175 | 0.995 | 0.452 | 0.803 |
| NA24385 | 29 | 1 | 20X | 176 | 4495 | 10054 | 0.983 | 0.447 | 0.807 |

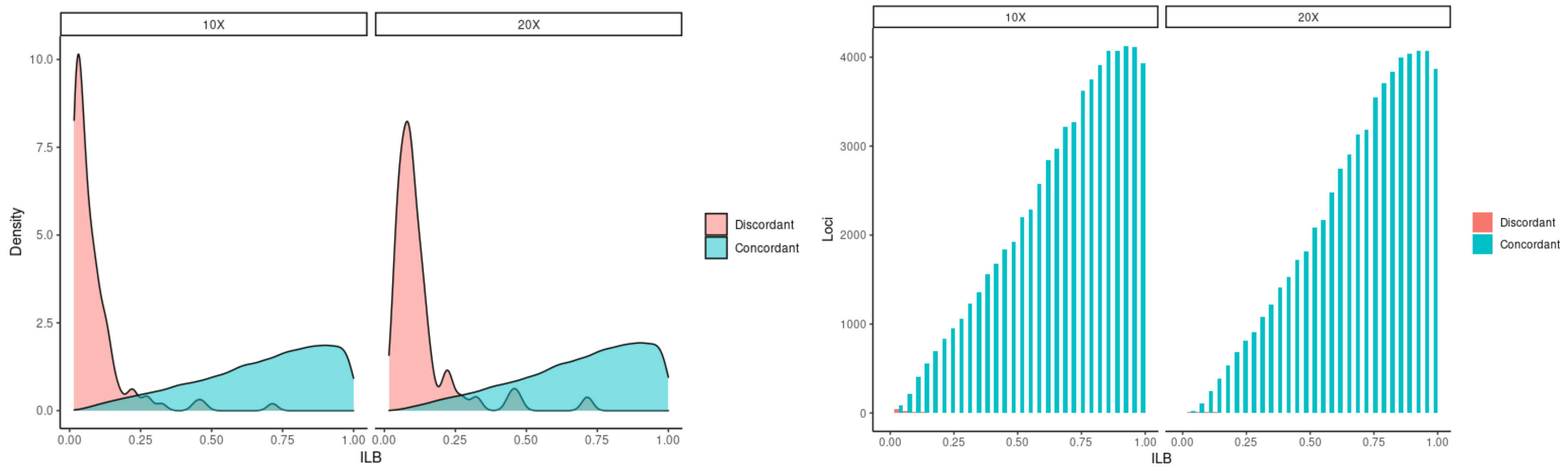


**Figure S5.** The density plot in A) displays the distribution of intralocus balance (ILB) values of true (concordant) and false (discordant) heterozygous calls, demonstrating that the majority of false heterozygous calls had low ILB. Based on this, we examined if, in addition to a read count threshold, an ILB threshold would be useful for assessing heterozygous genotypes. We applied a grid search approach using the F1 score to determine if there was an ILB threshold that would help eliminate these false calls. A range of thresholds was applied, where a genotype would be dropped if the ILB fell below the selected threshold. Each genotype was then categorized as the following: true positive (TP) – discordant calls that would be removed by the ILB threshold, false positive (FP) – concordant calls that would be removed by the ILB threshold, true negative (TN) – concordant calls that would be kept by the ILB threshold, or false negative (FN) – discordant calls that would be kept by the ILB threshold. These values were used to calculate recall (TP/(TP+FN)), precision (TP/(TP+FP)), and the F1 score (harmonic mean of precision and recall). The ILB threshold with the highest F1 (weighted for precision) was selected. Table S3 displays the ILB threshold values. However, applying the optimized thresholds only minimally removed discordant calls at the cost of removing many more concordant calls. This is in large part due to the significantly higher counts of concordant heterozygous calls (>60,000) than discordant heterozygous calls (<110) as displayed in the histogram in B). Note the discordant pink bars are barely visible.

**Table S3.** Optimized ILB thresholds specific to the 10X (0.06) and 20X (0.10) read count thresholds applied to the sensitivity study data. Due to the low impact the thresholds would have on removing false calls, an ILB threshold was not applied to the validation data.

| Read Count Threshold | ILB Threshold | Precision | Recall | Weighted F1 | Correctly Dropped (TP) | Incorrectly Dropped (FP) | Correctly Kept (TN) | Incorrectly Kept (FN) |
|---|---|---|---|---|---|---|---|---|
| 10X | 0.06 | 0.341 | 0.578 | 0.507 | 63 | 122 | 65253 | 46 |
| 20X | 0.10 | 0.120 | 0.604 | 0.334 | 32 | 235 | 62121 | 21 |

**Table S4.** The NA24385 libraries from the sensitivity study were additionally sequenced at the opposite plexity as was performed in that study to assess sequencing sensitivity. Thus, 0.05 and 0.1 ng samples were sequenced in a 4-plex pool, and 0.25 and 1.0 ng samples were sequenced in a 2-plex pool. The call rate, heterozygosity rate, and concordance rate mean values are presented here in comparison to the original sensitivity data for NA24385 (n=2 per condition). Call rate was calculated with all 10,230 SNPs across individual samples. Heterozygosity rate was calculated with called SNPs across individual samples. Concordance rate was calculated with 9,375 SNPs and 9,353 SNPs for NA24385 and NA24143, respectively, based on high confidence regions in the GIAB profiles, across individual samples.

| DNA Input (ng) | Analysis Threshold | Pool Plexity | Call Rate Mean | Heterozygosity Rate Mean | Concordance Rate Mean |
|---|---|---|---|---|---|
| 0.05 | 10X | 2 | 0.963 | 0.401 | 0.938 |
| 0.05 | 10X | 4 | 0.929 | 0.364 | 0.902 |
| 0.1 | 10X | 2 | 0.984 | 0.437 | 0.974 |
| 0.1 | 10X | 4 | 0.962 | 0.418 | 0.957 |
| 0.25 | 10X | 2 | 0.996 | 0.454 | 0.991 |
| 0.25 | 10X | 4 | 0.994 | 0.450 | 0.988 |
| 1 | 10X | 2 | 0.995 | 0.456 | 0.995 |
| 1 | 10X | 4 | 0.987 | 0.451 | 0.990 |

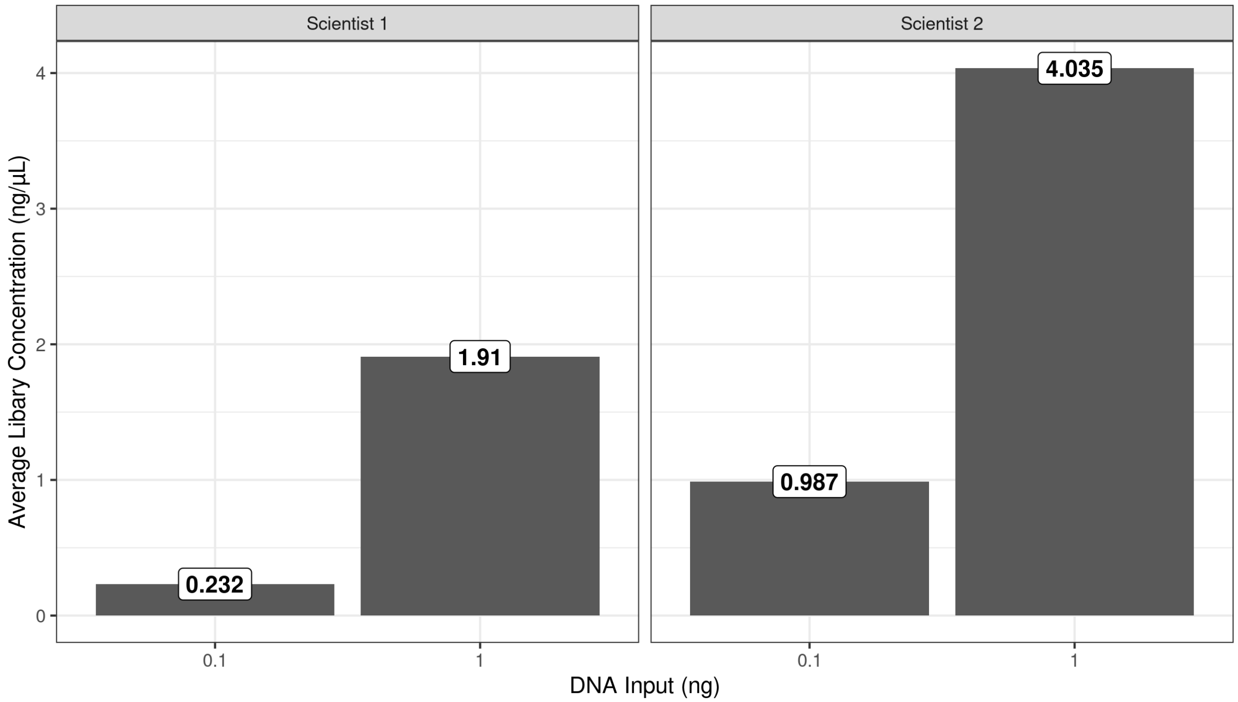


**Figure S6.** Average library concentration of 0.1 and 1.0 ng samples split by Scientist 1 and Scientist 2 replicates from the precision and accuracy study. The library concentration for Scientist 2 was 4.2 and 2.1X higher at inputs of 0.1 and 1.0 ng, respectively.

**Table S5.** Precision and accuracy study sample profile results, arranged by DNA input (ng). Scientist 1 results are taken from the sensitivity study and are duplicated here for ease of comparison.

| Scientist | Sample Name | Library ID | DNA Input (ng) | Analysis Threshold | Missed SNPs | Heterozygous SNPs | Called SNPs | Call Rate | Heterozygosity Rate | Intralocus Balance Mean |
|---|---|---|---|---|---|---|---|---|---|---|
| Scientist 1 | NA24143 | 30 | 0.1 | 10X | 250 | 4396 | 9980 | 0.976 | 0.440 | 0.627 |
| Scientist 1 | NA24143 | 32 | 0.1 | 10X | 204 | 4502 | 10026 | 0.980 | 0.449 | 0.635 |
| Scientist 2 | NA24143 | 73 | 0.1 | 10X | 135 | 4625 | 10095 | 0.987 | 0.458 | 0.653 |
| Scientist 2 | NA24143 | 75 | 0.1 | 10X | 155 | 4560 | 10075 | 0.985 | 0.453 | 0.657 |
| Scientist 1 | NA24385 | 21 | 0.1 | 10X | 136 | 4431 | 10094 | 0.987 | 0.439 | 0.602 |
| Scientist 1 | NA24385 | 23 | 0.1 | 10X | 186 | 4362 | 10044 | 0.982 | 0.434 | 0.597 |
| Scientist 2 | NA24385 | 69 | 0.1 | 10X | 89 | 4521 | 10141 | 0.991 | 0.446 | 0.618 |
| Scientist 2 | NA24385 | 78 | 0.1 | 10X | 61 | 4570 | 10169 | 0.994 | 0.449 | 0.608 |
| Scientist 1 | NA24143 | 31 | 1 | 10X | 157 | 4613 | 10073 | 0.985 | 0.458 | 0.813 |
| Scientist 1 | NA24143 | 38 | 1 | 10X | 99 | 4672 | 10131 | 0.990 | 0.461 | 0.817 |
| Scientist 2 | NA24143 | 76 | 1 | 10X | 148 | 4612 | 10082 | 0.986 | 0.457 | 0.821 |
| Scientist 2 | NA24143 | 77 | 1 | 10X | 106 | 4654 | 10124 | 0.990 | 0.460 | 0.825 |
| Scientist 1 | NA24385 | 22 | 1 | 10X | 220 | 4498 | 10010 | 0.978 | 0.449 | 0.786 |
| Scientist 1 | NA24385 | 29 | 1 | 10X | 55 | 4599 | 10175 | 0.995 | 0.452 | 0.803 |
| Scientist 2 | NA24385 | 70 | 1 | 10X | 47 | 4625 | 10183 | 0.995 | 0.454 | 0.807 |
| Scientist 2 | NA24385 | 72 | 1 | 10X | 55 | 4597 | 10175 | 0.995 | 0.452 | 0.803 |

**Table S6.** Mixture study sample profile results, arranged by mixture ratio and DNA input (ng).

| Mixture Ratio | Library ID | DNA Input (ng) | Analysis Threshold | Missed SNPs | Heterozygous SNPs | Called SNPs | Call Rate | Heterozygosity Rate | Intralocus Balance Mean |
|---|---|---|---|---|---|---|---|---|---|
| 1:2 | 58 | 0.1 | 10X | 199 | 7193 | 10031 | 0.981 | 0.717 | 0.466 |
| 1:2 | 59 | 0.1 | 10X | 243 | 7026 | 9987 | 0.976 | 0.704 | 0.465 |
| 1:2 | 56 | 1 | 10X | 42 | 8060 | 10188 | 0.996 | 0.791 | 0.499 |
| 1:2 | 57 | 1 | 10X | 7 | 8166 | 10223 | 0.999 | 0.799 | 0.500 |
| 1:5 | 62 | 0.1 | 10X | 312 | 6298 | 9918 | 0.970 | 0.635 | 0.470 |
| 1:5 | 63 | 0.1 | 10X | 328 | 6222 | 9902 | 0.968 | 0.628 | 0.478 |
| 1:5 | 60 | 1 | 10X | 22 | 7940 | 10208 | 0.998 | 0.778 | 0.505 |
| 1:5 | 61 | 1 | 10X | 31 | 7909 | 10199 | 0.997 | 0.775 | 0.502 |
| 1:20 | 64 | 1 | 10X | 78 | 6282 | 10152 | 0.992 | 0.619 | 0.611 |
| 1:20 | 65 | 1 | 10X | 50 | 6605 | 10180 | 0.995 | 0.649 | 0.592 |

**Table S7.** Negative control sample profile results, arranged by library ID. All SNPs called were homozygous, thus intralocus balance mean and heterozygosity rate are not reported.

| Study | Library ID | Sequencing Plexity | Analysis Threshold | Missed SNPs | Heterozygous SNPs | Called SNPs | Call Rate |
|---|---|---|---|---|---|---|---|
| Sequencing Sensitivity | 26 | 2 | 10X | 10198 | 0 | 32 | 0.003 |
| Sensitivity | 26 | 4 | 10X | 10217 | 0 | 13 | 0.001 |
| Sequencing Reproducibility | 26 | 4 | 10X | 10216 | 0 | 14 | 0.001 |
| Sequencing Sensitivity | 35 | 2 | 10X | 10215 | 0 | 15 | 0.001 |
| Sensitivity | 35 | 4 | 10X | 10223 | 0 | 7 | 0.001 |
| Sequencing Reproducibility | 35 | 4 | 10X | 10221 | 0 | 9 | 0.001 |
| Nonprobative | 55 | 4 | 10X | 10216 | 0 | 14 | 0.001 |
| Mixtures | 68 | 4 | 10X | 10208 | 0 | 22 | 0.002 |
| Precision and Accuracy | 71 | 4 | 10X | 10138 | 0 | 92 | 0.009 |
| Sequencing Reproducibility | 71 | 4 | 10X | 10154 | 0 | 76 | 0.007 |
| Precision and Accuracy | 74 | 4 | 10X | 10214 | 0 | 16 | 0.002 |
| Sequencing Reproducibility | 74 | 4 | 10X | 10214 | 0 | 16 | 0.002 |

**Table S8.** Nonprobative study sample profile results arranged by library ID.

| Sample Type | Donor ID | Sample Details | Library ID | Degradation Index | DNA Input (ng) | Analysis Threshold | Missed SNPs | Heterozygous SNPs | Called SNPs | Call Rate | Heterozygosity Rate | Intralocus Balance Mean |
|---|---|---|---|---|---|---|---|---|---|---|---|---|
| Bone | A | early surface decomp | 39 | 1.16 | 0.357 | 10X | 27 | 4784 | 10203 | 0.997 | 0.469 | 0.719 |
| Bone | A | early surface decomp | 40 | 1.22 | 1 | 10X | 183 | 4571 | 10047 | 0.982 | 0.455 | 0.740 |
| Bone | B | cremated | 41 | 2.67 | 1 | 10X | 787 | 3663 | 9443 | 0.923 | 0.388 | 0.553 |
| Bone | B | cremated | 42 | 2.67 | 0.1 | 10X | 3749 | 1126 | 6481 | 0.634 | 0.174 | 0.474 |
| Bone | C | burned | 43 | 9.92 | 1 | 10X | 219 | 4601 | 10011 | 0.979 | 0.460 | 0.724 |
| Bone | D | burned | 44 | 14.25 | 1 | 10X | 240 | 4651 | 9990 | 0.977 | 0.466 | 0.750 |
| Bone | D | burned | 45 | 14.25 | 0.1 | 10X | 727 | 3649 | 9503 | 0.929 | 0.384 | 0.541 |
| Bone | D | burned | 46 | 26.08 | 1 | 10X | 317 | 4636 | 9913 | 0.969 | 0.468 | 0.760 |
| Bone | E | embalmed | 47 | 3.69 | 1 | 10X | 31 | 4719 | 10199 | 0.997 | 0.463 | 0.708 |
| Bone | F | surface decomp | 48 | 32.40 | 0.444 | 10X | 2082 | 3154 | 8148 | 0.796 | 0.387 | 0.577 |
| Bone | G | buried | 49 | 4.24 | 0.271 | 10X | 196 | 4628 | 10034 | 0.981 | 0.461 | 0.602 |
| Shell casing | H | -- | 50 | -- | 0.052 | 10X | 917 | 3157 | 9313 | 0.910 | 0.339 | 0.525 |
| Shell casing | H | -- | 51 | -- | 0.062 | 10X | 1015 | 3150 | 9215 | 0.901 | 0.342 | 0.521 |
| Adhesive tape | NA24385 | -- | 52 | -- | 1 | 10X | 357 | 4399 | 9873 | 0.965 | 0.446 | 0.789 |
| Adhesive tape | NA24385 | -- | 53 | -- | 0.569 | 10X | 109 | 4574 | 10121 | 0.989 | 0.452 | 0.787 |
